# Selection for mitochondrial quality drives evolution of the germline

**DOI:** 10.1101/026252

**Authors:** Arunas L Radzvilavicius, Zena Hadjivasiliou, Nick Lane, Andrew Pomiankowski

## Abstract

The origin of the germline-soma distinction is a fundamental unsolved question. Plants and basal metazoans do not have a germline but generate gametes from somatic tissues (somatic gametogenesis), whereas most bilaterians sequester a germline. We develop an evolutionary model which shows that selection for mitochondrial quality drives germline evolution. In organisms with low mitochondrial mutation rates, segregation of mutations over multiple cell divisions generates variation, allowing selection to optimize gamete quality through somatic gametogenesis. Higher mutation rates promote early germline sequestration. Oogamy reduces mitochondrial segregation in early development, improving adult fitness by restricting variation between tissues, but also limiting variation between early-sequestered oocytes, undermining gamete quality. Oocyte variation is restored through proliferation and random culling (atresia) of precursor cells. We predict a novel pathway from basal metazoans lacking a germline to active bilaterians with early sequestration of large oocytes subject to atresia, allowing the emergence of complex developmental processes.

## Introduction

In distinguishing between the germline and soma, Weismann argued that the division of labour enabled the specialization of cells in somatic tissues, permitting greater organismal complexity^1^. In contrast, germline cells alone retain the capacity to provide genetic information for future generations, and never form somatic cells^2^. Without the specialization enabled by the germline, complex multicellular animals with post-mitotic tissues such as brain might be impossible. But the division of labour cannot account for the origin of the germline, as all plants and many animals (including tunicates, flatworms and Cnidaria^3^) have differentiated tissues but do not sequester a germline, instead generating gametes from pluripotent stem cells in somatic tissues (somatic gametogenesis).

The best known hypothesis for the evolution of the germline relates to selfish competition between the cells of an individual: by resolving evolutionary conflict, the strict distinction between germline and soma stabilises multicellular cooperation^4-6^. According to this theory, plants did not evolve a germline because their rigid cell walls restrict cell movement, limiting the systemic effects of any parasitic cell lines^4^. In contrast, animal cells lack a rigid wall, making them more vulnerable to parasitic cell lines that undermine organismal function^4^. Sequestering a germline theoretically limits this competition. However, because cells in multicellular organisms normally derive from a single cell (unitary development) new selfish mutations must arise within a single generation; if these mutants are inherited then all cells in the offspring will carry the selfish mutation, so there are no longer any non-selfish cells to exploit^7,8^. Conflict can therefore only emerge when there is a high mutation rate and a large number of cell divisions, which is true of most metazoans^7^. But that does not explain why many metazoan groups do not sequester a germline, or why some phyla (such as Ectoprocta and Entoprocta secondarily returned to somatic gametogenesis^9^.

A second line of thinking dating back to Weismann emphasises the protected environment of a sequestered germline^1,10^. By restricting mutations in gametes, the germline enhances germ-cell quality, contrasting with the ‘disposability’ of the soma^11^. Avoiding the accumulation of mutations in nuclear genes associated with greater metabolic work can theoretically favour differentiation of a germline^12^. But testes have a high metabolic rate and sperm are produced continuously through life, with 30 cell divisions by puberty and 400 divisions at the age of 30 in humans, contributing a large number of new mutations to offspring^13^. From this point of view, the male germline is equivalent to tissues such as the bone marrow, merely specialising in the mass production of cells of a particular type.

Focusing on the origin of germline sequestration therefore shifts the problem specifically to female gametogenesis. Female gametes are sequestered early in development in a transcriptionally repressed state, with meiosis arrested in prophase I and mitochondria in a state of functional quiescence^14^. Strikingly, female gametes (oocytes) are usually large cells packed with mitochondria, with as many as 10^6^ copies of mitochondrial DNA in mammalian oocytes^15^. Mitochondrial DNA is usually inherited uniparentally: male mitochondria are either excluded from the zygote or destroyed on entry to the oocyte^16^. These traits point to mitochondrial function as being central to germline evolution. The possibility that selection for mitochondrial quality could have driven the evolution of the germline has been raised before^14,17,18^, but never formally addressed. We show that selection for mitochondrial quality can explain many long-puzzling features of germline evolution.

## Model of germline evolution

We consider the evolution of the germline in a model multicellular organism with variable tissue-level differentiation. Gametes are formed either from adult somatic tissues or early in embryogenesis (Fig. 1a & b). The quality of gametes and fitness of the organism depends on the number of mitochondrial mutations. These build up due to copying errors in mitochondrial genes at a rate *µ*_S_ per cell division (Fig. 1c). Early sequestration of a germline restricts the number of cell divisions to gamete production, and therefore the number of copying errors compared with gametes differentiated from adult tissues. Mutations can also result from ‘background’ damage, *µ*_B_, per unit time (caused by oxidative damage or UV radiation; Fig. 1d), as cells can accumulate mitochondrial mutations even when not actively dividing^19-22^. So background mutations affect gametes whether they are derived from an early germline or later from somatic cells.

**Figure 1.**
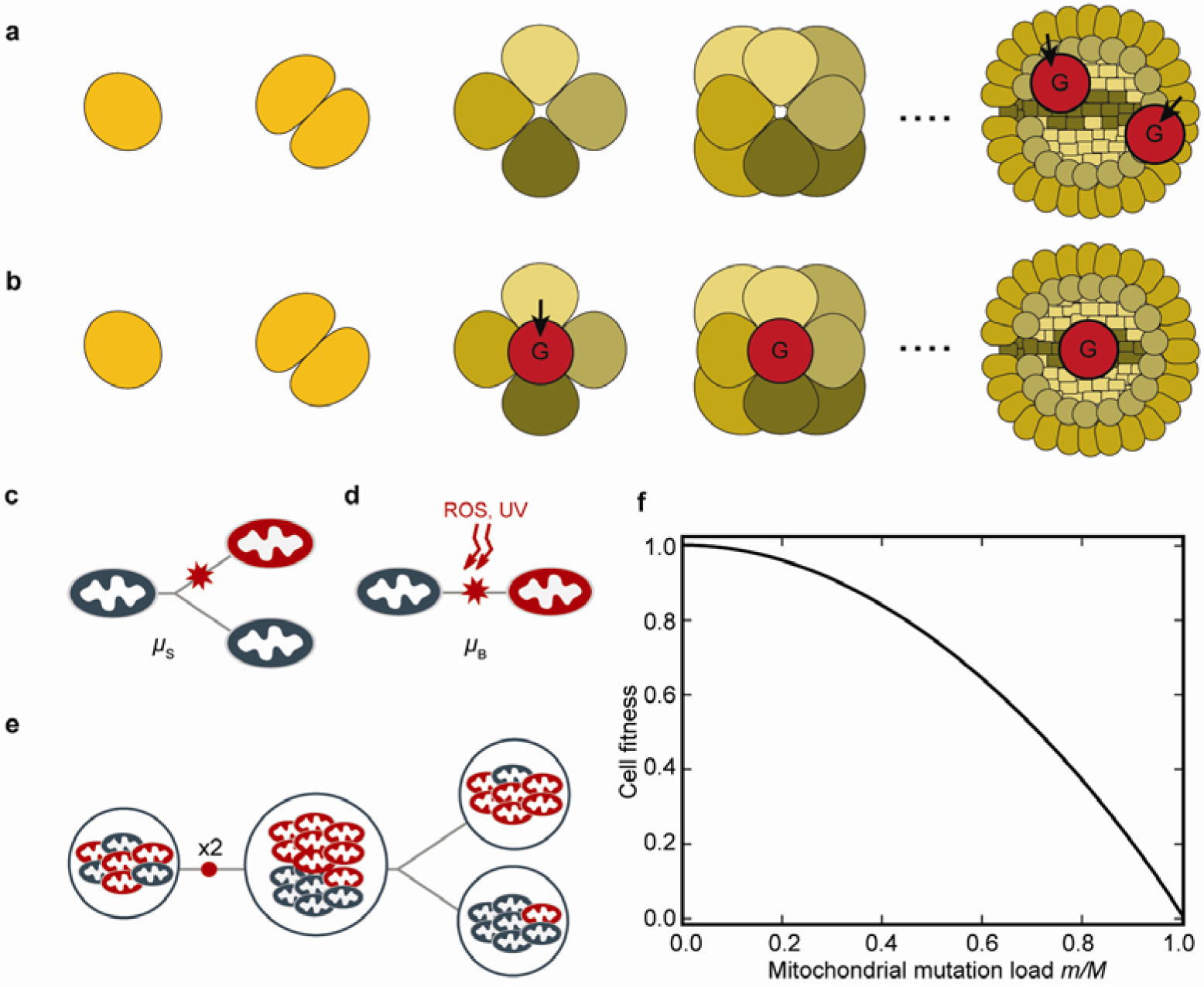
Life cycle of the model multicellular organism (a) Development from zygote (left-hand side) to adult, showing early tissue differentiation (cells with differing shade) and late formation of gametes (G) from somatic cells in the adult (somatic gametogenesis). **(b)** Equivalent multicellular development depicting sequestration of gametes early in development (early germline). Dotted line indicates further development; adult and gamete cells not drawn to scale. **(c)** Copying errors (*µ*_S_) during replication of mitochondrial DNA. **(d)** Mutations caused by background damage (*µ*_B_) from e.g. ultraviolet radiation (UV) or reactive oxygen species (ROS). **(e)** Doubling followed by random segregation of mitochondrial mutants (red) at cell division increases variance between daughter cells. **(f)** Concave fitness function, in which cell fitness declines non-linearly with the accumulation of mitochondrial mutations (*µ*_S_ + *µ*_B_) as seen in mitochondrial diseases^23^.

Unlike nuclear genes, which are clonally transmitted through mitosis, the mitochondrial population doubles and segregates at random to daughter cells at each cell division. Some daughter cells receive more, others fewer, mitochondrial mutations (Fig. 1e). With somatic gametogenesis (i.e. late differentiation of germ cells), segregational variation has the capacity to generate gametes that carry few mutants or are even mutation free. The more rounds of cell division, the greater the degree of segregation, the higher the variance in mutation load, and the larger the number of gametes carrying very few or no mitochondrial mutations (Fig. 2a). The effect of segregation is dampened with larger numbers of mitochondria per cell, as this diminishes drift^15^ (Fig. 2b). At higher copying-error rates (high *µ*_S_), segregation still generates increased variance between daughter cells, but the proportion of gametes with few or no mutations falls to nearly zero, making segregation and selection less effective at maintaining mitochondrial quality (Fig. 2c). Including selection at the level of mitochondria equates to a change in the mutation rate. For example, removing deleterious mitochondria by mitophagy effectively reduces the mutation rate, whereas faster replication of selfish or impaired mitochondria increases the mutation rate. For simplicity we therefore ignore selection at the level of mitochondria. Likewise, although the severity of mitochondrial mutations is variable, we fix each to have a similar effect for ease of analysis.

**Figure 2.**
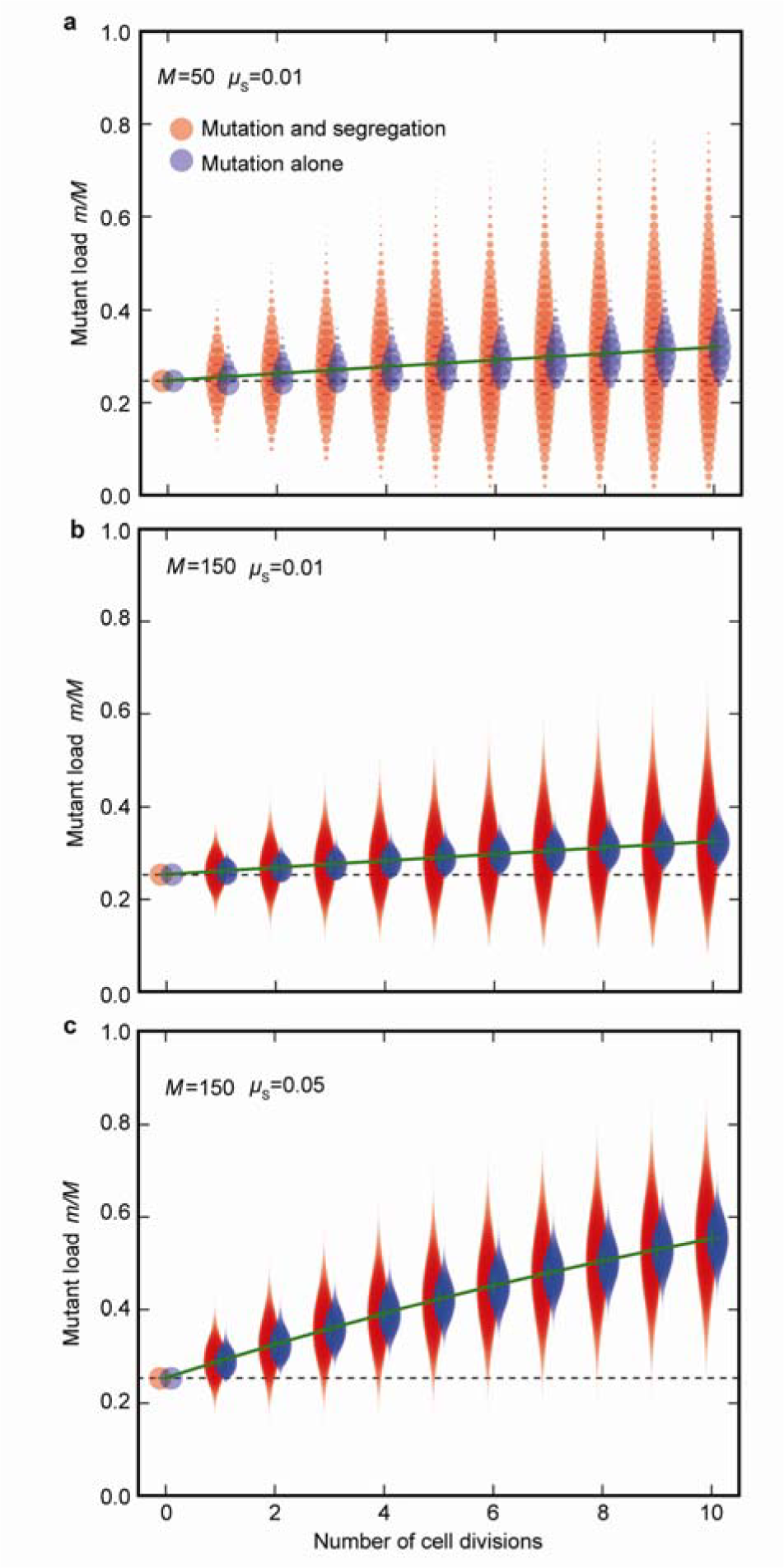
Segregation of mitochondrial mutants at cell division generates variance. At each cell division, the variance in mitochondrial mutant load (*m*/*M*) between daughter cells (red plots) increases due to mutational input as well as random segregation, generating cells with both more (above dotted line) and fewer (below dotted line) mutations than the zygotic cell. The blue plots show the variance in mutant load at each division due to mutation alone, without any random segregation. **(a)** The mutant load increases slowly with each cell division (green line) when the rate of copying errors is low (*µ*_S_ = 0.01). **(b)** Increasing mitochondrial number (*M*) decreases the variance in mutant load per cell (red plots) as the effects of random segregational drift are dampened. Fewer high-quality daughter cells lacking mitochondrial mutations are now generated. **(c)** Increasing copying errors (*µ*_S_ = 0.05) increases the mutant load in daughter cells; segregation no longer generates many high-quality daughter cells with fewer mutations than the zygote. Data were derived from a starting number of mutants in zygotes *m*= 0.24 *M*, and then run iteratively through successive cell divisions as described in the first section of Methods.

Overall, the resulting mitochondrial population in somatic cells is subject to a mutational load, which impacts on cell fitness

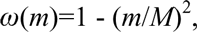

where *m* is the number of mutant mitochondria and *M* is the total size of the mitochondrial population in each cell. This fitness function is concave, and assumes that a large number of mutants must accumulate before cell function is significantly undermined (Fig. 1f). This is the case for mitochondrial diseases, where mutant load can reach 60–80% before symptoms are detectable^23^. Other possible fitness functions (e.g. linear or convex) do not affect the general outcome of our modelling but produce slightly different thresholds for germline evolution. We therefore concentrate on a concave fitness function as this is the most empirically justifiable.

Another important dimension of fitness in multicellular organisms is the number and quality of tissues, and their mutual interdependence. We take the fitness of a somatic tissue to be the mean fitness of its constituent cells, each of which reflects the accumulation and segregation of mutations as described above. We assume strong negative epistasis between tissues, so that adult fitness is determined by the quality of the worst tissue. If by chance a disproportionate number of mitochondrial mutations segregate into precursors of one tissue, the fitness of the adult is severely impaired; conversely if mutations are distributed equally into the precursor cells of all tissues, organismal fitness is improved, as each tissue would have equivalent quality. This corresponds to bilaterians in which tissue-precursor cells are defined early in embryonic development, and organ function is mutually interdependent^4^. Failure of one organ (e.g. heart, liver) causes death or debilitation of the organism. In contrast, ancestral metazoans such as sponges have different cell types, but do not differentiate distinct tissue precursors early in development^24^, so there is unlikely to be significant tissue-specific epistasis effects on adult fitness. Plant tissues fall between sponges and bilaterians. Nascent tissue organization in seeds is equivalent to the embryonic organization of metazoans^25^, but the repetitive modularity of adult tissues means that organ failure due to accumulation of mitochondrial mutations (e.g. of a flower, leaf) has a minimal effect on the fitness of the organism. In the model, plants are regarded as having fewer mutually dependent tissues compared with bilaterians, as there is less negative epistasis between the component parts.

Adult fitness thus depends on the number of mitochondrial mutations inherited by the zygote, the number of new mutations arising during development, how these mutations segregate into cells and tissues, and the degree of tissue-level epistasis. As we will show, this combination determines whether it is better to sequester germ cells early in development in a germline, or derive them from adult somatic cells.

## Germline evolution depends on mitochondrial mutation rate

The main benefit of early sequestration of germ cells is to reduce the number of cell divisions before gamete production, so reducing the net input of copying errors (*µ*_S_) in gametes. An early germline raises the mean fitness of offspring in the next generation, but comes at the cost of reduced segregational variance. When gametes are derived later in development, there is more chance for segregation to generate gametes with lower numbers of mitochondrial mutations, which facilitates selection in the offspring, improving fitness over generations (Fig. 2). The tension between these two forces determines the advantage (or disadvantage) of a germline.

When mitochondrial mutation input through copying errors is low, the benefit of increased variance between gametes tends to outweigh the benefit of sequestering an early germline (Fig. 3a). In other words, at low *µ*_S_ the benefit of curtailing cell division in an early-sequestered germline is less than the benefit of increased variance between gametes (forming some gametes with few or no mutations) generated by somatic gametogenesis (Fig. 2). Note that if copying errors were the only form of mutation, then selection would always favour an early germline (Fig 3a bottom left corner). However, oocytes sequestered early in development continue to accumulate mutations as a result of background damage (*µ*_B_)^19^. These mutations can only be segregated out through further rounds of cell division, which favours late somatic gametogenesis. Germline sequestration is therefore unlikely to evolve in organisms with low *µ*_S_ but relatively high *µ*_B_ (Fig 3a, bottom right corner), such as those that have long lifespan or live in damaging environments (e.g. high UV exposure). These conditions are true for basal metazoans and plants. Unlike most animal mitochondrial genomes, basal metazoans including sponges, corals and placozoans all have very low copying error rates^26-28^ (low *µ*_S_) and long lives (high *µ*_B_) which readily explains why these major phyla lack a germline. Likewise, most plants have low *µ*_S_ (10-20 fold lower than in their nuclear genomes^29^, and 50-100 times lower than typical animal mitochondrial genomes^30^) while being exposed to high UV (high *µ*_B_) in their phototrophic niche.

**Figure 3.**
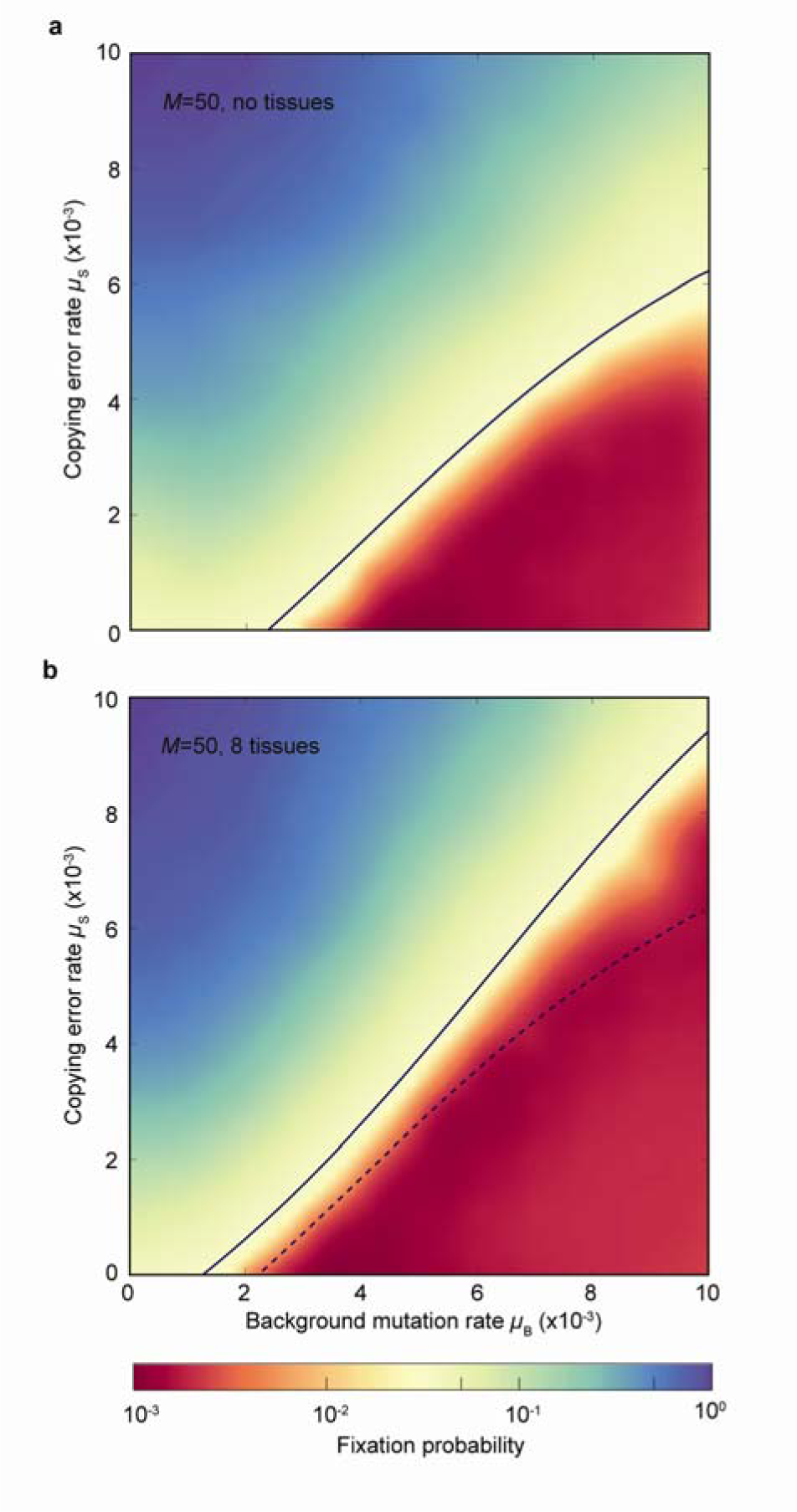
Germline evolution depends on mitochondrial mutation rate (a) Heat map showing fixation probability of an allele encoding early germline sequestration (introduced at a frequency of 0.05, see Methods) in a simple organism that lacks tissue differentiation, in relation to the rate of copying-errors (*µ*_S_) and background damage (*µ*_B_). Early germline sequestration is favoured by higher *µ*_S_ and lower *µ*_B_ (blue, top left). The early germline allele is selected against in organisms with low *µ*_S_ and high *µ*_B_ (red, bottom right), conditions that instead favour somatic gametogenesis. The solid line represents neutrality. **(b)** Increasing the number of tissues to 8 makes it harder to fix an early germline – the region shaded in red expands (solid line versus dotted line) so germline fixation now requires higher *µ*_S_ and lower *µ*_B_ compared with (a).

Importantly, increasing the number of tissues (Fig. 3b) or mitochondria (Extended Data Fig. 1), which both correspond to greater complexity, does not enhance the likelihood of germline evolution. The reason again relates to segregation. In organisms with a single tissue, mean adult fitness closely follows the mutation load in the zygote, with some variation induced by mutation accumulation (Fig. 4a). In contrast, with multiple tissues, mutations segregate differentially into distinct tissues. As the fitness of an organism depends on the mutational state of the worst tissue, this magnifies the fitness disadvantage of an inherited mutation load. So there is a substantial increase in variance and reduction in mean adult fitness relative to the inherited zygote fitness (Fig. 4b). This gives an added advantage to the late production of gametes (Fig. 3b), as the extra segregational drift increases the variance in mutation load between gametes. Higher variance between individuals with somatic gametogenesis leads to higher average adult fitness, compared to competitors with early germline sequestration. Increasing mitochondrial number (larger *M*) also reduces the variance in mutation load between tissue precursor cells, which likewise improves adult fitness (Fig. 4c). This again militates against the production of early germline sequestration with less segregational variation produced by fewer cell divisions (Extended Data Fig. 1). Overall, then, multiple tissues and high numbers of mitochondria do not drive the evolution of early germline sequestration, but rather reinforce the stability of somatic gametogenesis in plants and many metazoans.

**Figure 4.**
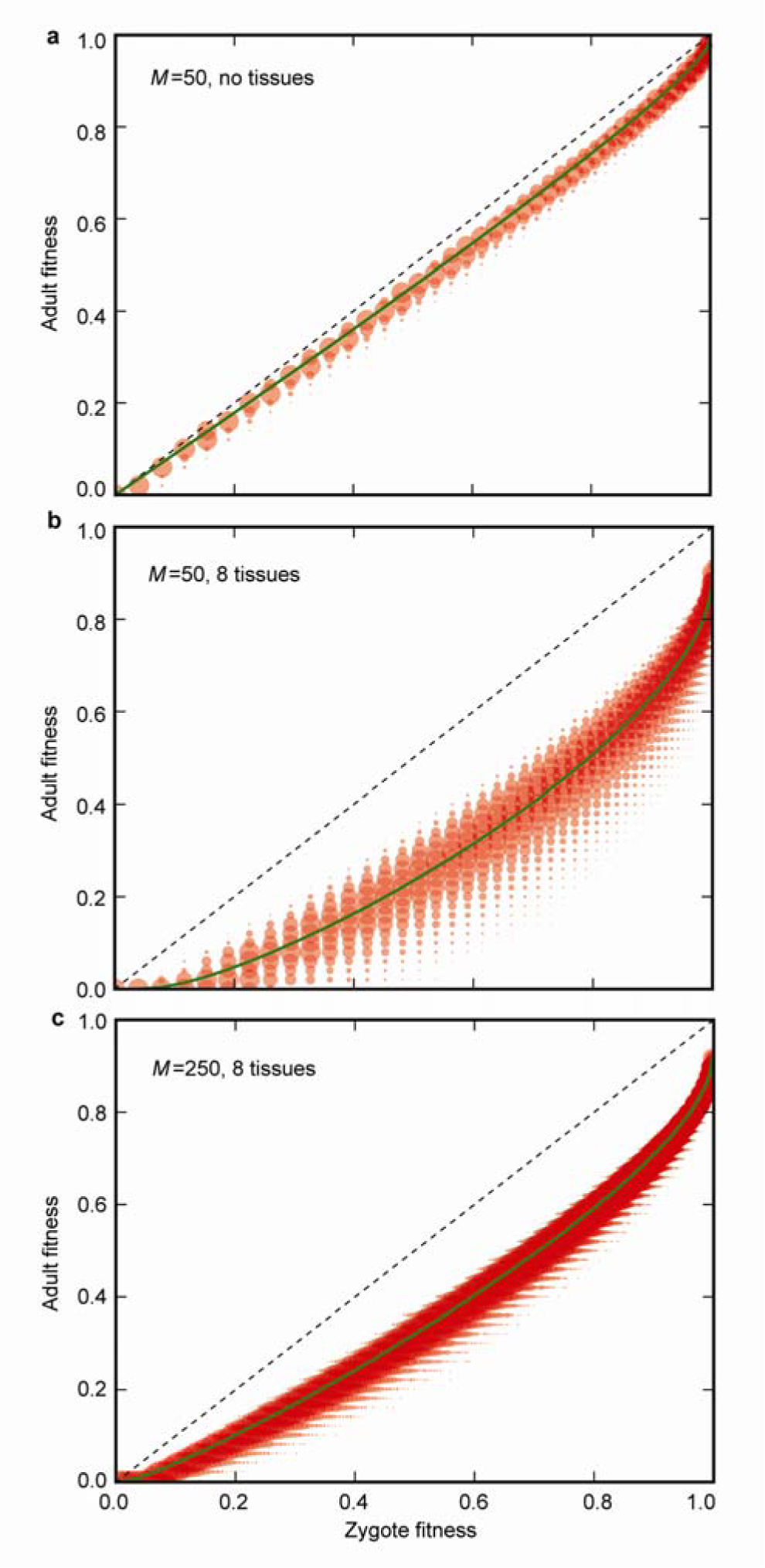
Mitochondrial segregation undermines adult fitness. Adult fitness is a function of zygote fitness (the mutation load inherited), mutational input and random segregation during development. **(a)** In organisms with no tissue differentiation adult fitness is similar to zygote fitness, as the number of new mutations accumulating within a single generation is limited, and variance in mutant load between cells within a tissue has no effect on adult fitness. **(b)** In organisms with early differentiation of multiple tissues, adult fitness is undermined by segregational variance, as some tissue-precursor cells receive a higher mutant load than others, and adult fitness depends on the function of the worst tissue. **(c)** Increasing the number of mitochondria decreases the variance in mutant load between tissue-precursor cells, and so reduces the loss in adult fitness caused by random segregation. Parameter values *µ*_S_ = 0.01, *µ*_B_ = 0.005, 10 cell divisions to adulthood, and a lifespan equivalent to 40 cell division cycles.

The one factor that reliably drives sequestration of an early germline is an increase in the rate of mitochondrial copying errors, *µ*_S_ (Fig. 3a, top left). With high *µ*_S_, it is better to avoid mutation accumulation and move gamete production into an early sequestered germline, as segregation is no longer strong enough to generate fitter gametes with few or no mitochondrial mutants (Fig. 2c). There are two animal groups in which early germline sequestration is widespread – bilaterians^4,9^ and ctenopohores^31^. In line with the prediction of our model, both groups have high mitochondrial mutation rates, 10-50 times faster than their mean nuclear mutation rate^32,33^, and these mutations are largely produced by copying errors^34^. Why did the mitochondrial mutation rate increase in the lineages giving rise to bilaterians and ctenophores? Increasing oxygen levels in the late Neoproterozoic^35^ enabled predation for the first time, because energy conservation from aerobic respiration approaches 40% compared with < 10% for fermentation, making predation virtually impossible in anoxic worlds^36,37^. In the early Cambrian, predation drove an evolutionary arms race leading to greater size and physical activity in bilaterians and probably ctenophores^38^. Greater activity increased rates of tissue turnover, protein synthesis and replication of mitochondrial genes, contributing to an elevated *µS* relative to sessile early metazoans^39^. A link between mitochondrial mutation rate and germline evolution is corroborated by the Ceriantharia, which unlike other Anthozoa have fast-evolving mitochondrial DNA^40^, suggesting a relatively high *µ*_S_; strikingly, their larvae have gonads and are apparently paedogenetic^41^, implying the evolution of germline sequestration in this group. Conversely, the secondary evolution of somatic gametogenesis in Ectoprocta and Entoprocta^9^ may have been favoured by a falling *µ*_S_. Little is known about mitochondrial sequence divergence in these groups, but we predict that *µ*_S_ will be lower in these groups than in related metazoans that retain a germline.

## Oogamy improves adult fitness and gamete quality

Oogamy has typically been interpreted as an outcome of disruptive selection for gamete specialization, with large eggs for provisioning, contrasting with numerous small sperm for success in fertilization^42^. Oogamy is universal in extant metazoan groups and common to multicellular organisms in general, both with and without germline, but is most extreme in complex organisms with a germline^43^. Our analysis specifies more precisely the benefits that flow from the generation of large oocytes with high (or low) mitochondrial numbers and the relationship of this to the evolution of a germline.

Oogamy offers a temporary increase in mitochondrial number (*M*) that feeds through to the zygote. Imagine that oocytes undergo *Q* additional rounds of mitochondrial replication without cell division, producing large oocytes with 2*^Q^M* mitochondria. After fertilization, the first few rounds of cell division partition the initial mitochondrial population in the zygote between the daughter cells, without the need for further replication, until the baseline *M* mitochondria is restored (Fig. 5a). Note that a large oocyte does not alter the total input of mitochondrial mutations per generation, because replication events lost from early development are simply moved later into gametogenesis. Nor does it impact on the net input of background mutations (*µ*_B_), which is simply a function of lifespan unchanged by oogamy.

**Figure 5.**
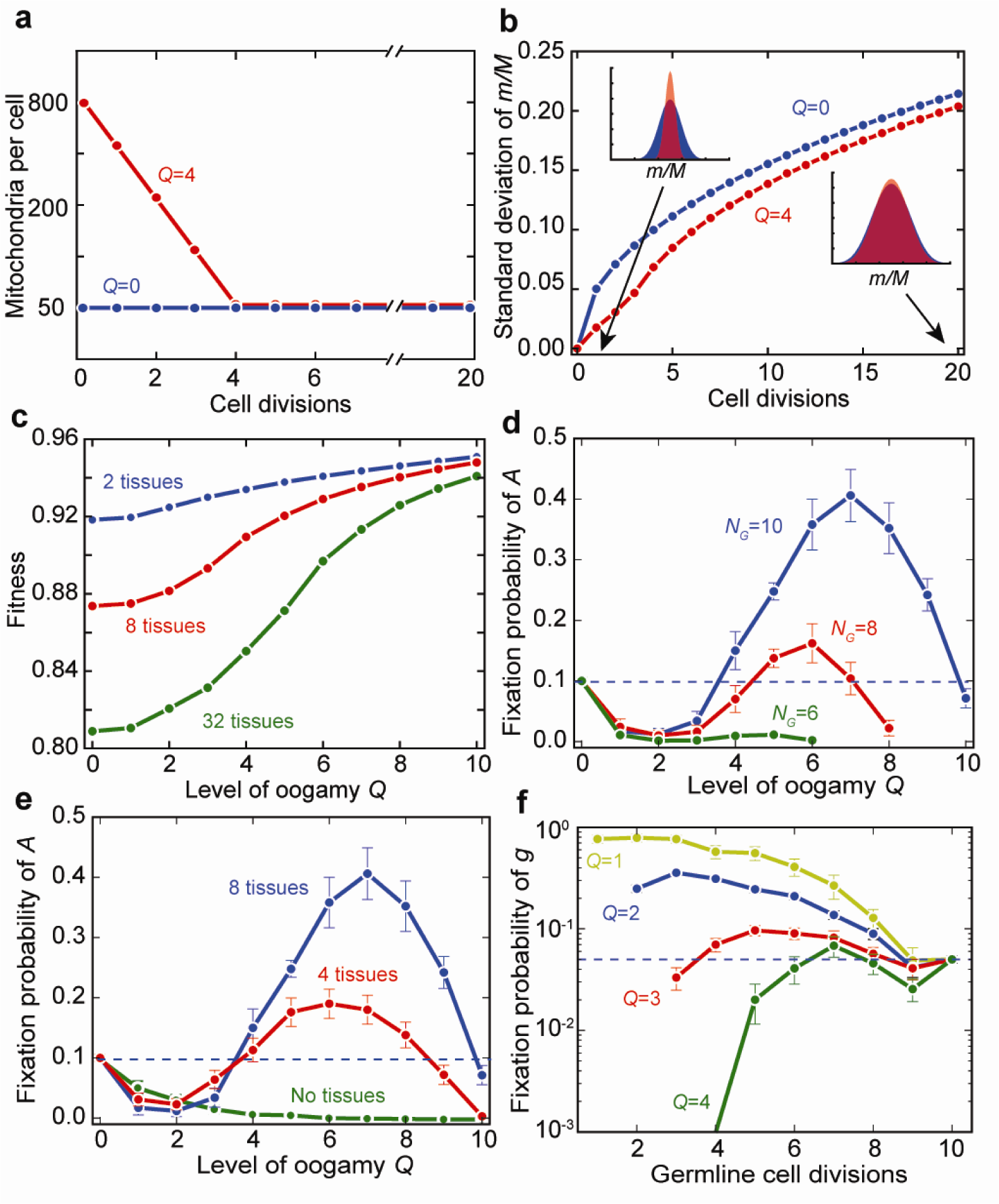
Oogamy improves adult fitness by temporarily suppressing variance (a) The mitochondria in the zygote are partitioned into daughter cells at each cell division. Under isogamy (*Q* = 0), the zygote contains the same number *M* mitochondria as normal somatic cells. In contrast, with oogamy the zygote contains a much larger number of mitochondria which are partitioned without further replication until the standard mitochondrial number (*M*) is restored. For example, when *M* = 50 and *Q* = 4, the zygote has 2*^Q^M* = 800 mitochondria, which are partitioned to daughter cells over 4 rounds of cell division to restore *M*. **(b)** A large oocyte (*Q* = 4, red) suppresses the variance in mutation load (*m*/*M*) in the first few rounds of cell division (left inset) compared with a small oocyte (*Q* = 0, blue). This early difference in variance is virtually lost after 20 rounds of cell division (right inset). The initial number of mitochondrial mutants in the zygote is 2*^Q^M*/2, and the segregation is modelled as described in the Methods without the further accumulation of mutations. **(c)** The early reduction in variance produced by oogamy improves adult fitness in organisms with multiple tissues. The more tissues, the greater *Q* needs to be to offset the loss of adult fitness. The initial mutant load in the zygote is 2*^Q^M/*5, *µ*_S_ = 0.01, *µ*_B_ = 0.005. **(d)** The fixation probability of an allele *A* specifying oogamy *Q* depends on the timing of gamete differentiation (*N_G_*). The allele *A* fixes with somatic gametogenesis (*N_G_* = 10) as more rounds of cell division generates greater variance between gametes, improving fitness over generations. But *A* does not fix with early germline sequestration (*N_G_* = 6 or less), as fewer rounds of cell division do not generate enough variance between gametes for selection to efficiently eliminate mutants. Mutation rates are set to *µ*_S_ = 0.01, *µ*_B_ = 0.005. **(e)** With somatic gametogenesis (*N_G_* = 10) the allele *A* encoding oogamy *Q* fixes more readily with a larger number of tissues and should not fix in organisms lacking tissues (but see Extended Data Fig. 3). Parameter values *µ*_S_ = 0.01, *µ*_B_ = 0.005. **(f)** The forces of mutation and segregation conflict in organisms with multiple tissues, where high *Q* improves adult fitness but high *µ*_S_ forces early germline sequestration. An allele *g* that specifies the number of cell divisions before gamete formation favours an intermediate number of cell divisions, in which mutation accumulation and segregational variance are balanced depending on *Q*. Parameter values *µ*_S_ = 0.02, *µ*_B_ = 0.001. In all panels, *M* = 50.

The temporary increase in *M* profoundly dampens the opportunity for segregational variance to build up in the early cell divisions (Fig. 5b, left-hand side), which greatly reduces the risk of one tissue inheriting a disproportionate number of mitochondrial mutants. This early suppression of variance is most valuable in organisms with multiple tissues, where oogamy greatly improves adult fitness (Fig. 5c). That in turn allows oogamy to readily fix, at least in organisms with somatic gametogenesis (Fig. 5d, *N*_G_ = 10). Once the early divisions of the zygote have allowed cells to regain the standard low level of *M* mitochondria, the many subsequent cell divisions before the end of development allow plenty of opportunity for segregational variation to generate fitter gametes with few or no mitochondrial mutants (Fig. 5b, right-hand side).

Strikingly, plants and basal metazoans have smaller numbers of mitochondria in oocytes (i.e. lower values of *Q*) than are typical of bilaterians^44-46^, suggesting that ‘mitochondrial oogamy’ is distinct from the large size of oocytes for provisioning. This lower degree of mitochondrial oogamy can be explained by the lower mutual dependence of adult fitness on organ function in plants (i.e. less negative epistasis between tissue fitness). In the model this is equivalent to a smaller number of tissues. Fig. 5e shows that the optimal *Q* is indeed lower with fewer tissues (e.g. 4 compared with 8 tissues). Nonetheless, even in plants there is a degree of embryonic tissue determination in early development (e.g. in seeds^25^), so moderate mitochondrial oogamy is predicted. In the case of basal metazoans that lack tissue organization such as sponges, the model suggests that oogamy would not be favoured (Fig. 5e, no tissues). However, other factors, in particular the degree of uniparental inheritance of mitochondria, can still favour the spread of oogamy as well as spermatogenesis – the production of small male gametes (Extended Data Fig. 2).

In contrast to somatic gametogenesis, the model analysis indicates that oogamy becomes less advantageous as the germline moves earlier in development (Fig. 5d, *N*_G_ = 6). Oogamy has the down-side that reduced segregation early in development carries over into gametes when they are drawn from an early germline. This reduces variation and the production of gametes with few or no mitochondrial mutations, which keenly weakens the response to selection. This disadvantage comes to outweigh the advantage gained by raising average tissue fitness through the suppression of segregation in early embryos. Yet the observation is that active bilaterians invariably combine early germline sequestration with large oocytes, in the extreme packed with several orders of magnitude more mitochondria than are present in normal cells (10^6^ vs. ~10^3^)^14,15^. So the model seems at odds with observations. In short, in active bilaterians with differentiated, mutually dependent tissues and high *µ*_S_, there is strong selection for early germline sequestration to limit the accumulation of mutations in gametes (Fig. 4b, top left), but equally strong selection for oogamy to minimize variation between tissue precursor cells (Fig. 5d), with the two forces opposing each other.

The solution to this paradox is suppression of the accumulation of copying errors (*µ*_S_), while maximising the variance between oocytes via segregation during germline development. In human development > 6 million oogonia are generated by mitotic proliferation in the first half of foetal development, followed by seemingly random apoptotic death of all but 500,000 of them (i.e. > 90%) by the start of puberty (atresia)^47^. A similar pattern is common across most bilaterian metazoans^48^. The proliferation of oogonia helps restore segregational variance among female gametes, which now have the chance of carrying few or no mitochondrial mutations (Extended Data Fig. 3). This “over supply” is rectified by eliminating 9 out of every 10 oogonia, the survivors being ‘islands’ of segregational variance, more different to each other than would be the case if fewer divisions were used to generate an equal number of oocytes. The balance between restricting *µ*_S_ through early germline sequestration and increasing variance through oocyte proliferation defines the number of cell divisions before atresia (Fig. 5f) – enough to generate variance, but less than the hundreds of cell divisions in the male germline or somatic gametogenesis. Thus increasing *Q* favours germline sequestration at an intermediate stage, neither early nor late, but after some rounds of cell division to generate variance between gametes (Fig. 5f).

Restoring variance necessarily generates new mutations via copying errors (*µ*_S_), but the total build-up of new mutations can be lowered by adaptations that suppress the accumulation of background damage (*µ*_B_) in female germ cells. These adaptations include protecting oocytes in an internal ovary, provisioning them from follicular cells, and repressing the transcription of mitochondrial genes and active respiration, which can produce reactive oxygen species^14,17,18^. Accordingly, early sequestration of large oocytes is most favoured when accompanied by specific lowering of *µ*_B_ (Extended Data Fig. 4), giving rise to quiescent oocytes. This is the case in the female germline of virtually all bilaterians^14^.

## Conclusions

For the last thirty years, the dominant explanation for the evolution of the germline has been that it was essential for the emergence of multicellular organisms through the suppression of selfish conflict between the cells that make up an individual. At its heart, this viewpoint lacks a rationale for the evolutionary stability of numerous organisms that lack a dedicated germline. It also sees the restricted mutational environment of the female germline as of secondary importance.

Here we have turned this paradigm on its head, locating the key driving force in selection for mitochondrial quality. This reflects the interplay of mutation and segregation of mitochondrial mutations in gametes and tissues. With low copying-error rates (*µ*_S_), the combination of segregation and selection improves mitochondrial quality over generations, favouring somatic gametogenesis in plants and simple metazoans that lack multiple tissues or high numbers of mitochondria. Higher *µ*_S_ drives early germline sequestration, which ultimately permits greater tissue differentiation and complexity in groups such as bilaterians and ctenophores. The increase in *µ*_S_ probably related to the rise in oxygen shortly before the Cambrian explosion^37^. We find that the evolution of complex development, with strong negative epistasis between tissues, requires suppression of mitochondrial variance between tissue-progenitor cells. This is achieved by large oocytes containing high mitochondrial numbers. That in turn risks the loss of segregational variance between oocytes sequestered in the germline early in development. The solution was to reintroduce segregational variance through the mitotic proliferation of oogonia, followed by germline atresia, maximizing the variance between the surviving oocytes.

These findings set out a completely novel hypothesis for the origin and evolution of germline sequestration, which predicts the traits of groups from plants and sponges to ctenophores and bilaterians. Unlike other hypotheses, selection for mitochondrial quality can account for the stability of both somatic gametogenesis and early germline sequestration. The requirement for segregational variation in the germline also elucidates the potential risks associated with mitochondrial heteroplasmy^16^ and the need for strict uniparental inheritance in more active animals^52^. The unusual population genetics of mitochondrial mutation and segregation can uniquely explain the evolution of the germline in complex bilaterians.

## Acknowledgments

The authors acknowledge the use of the UCL Legion High Performance Computing Facility (Legion@UCL), and associated support services, in the completion of this work. A.L.R. is supported by a CoMPLEX PhD studentship from the Engineering and Physical Sciences Research Council, Z.H. by an EPSRC Research Fellowship (EP/L504889/1), N.L. by the Leverhulme Trust (RPG-425) and the Provost’s Venture Research Prize, and A.P. by grants from the Natural Environment Research Council (NE/G00563X/1) and the Engineering and Physical Sciences Research Council (EP/F500351/1, EP/I017909/1, EP/K038656/1).

### Author Contributions

All authors designed the study, A.L.R., N.L. and A.P. analyzed the results and wrote the paper. A.L.R. carried out the programming and simulations.

---

**Extended Data Figure 1.**
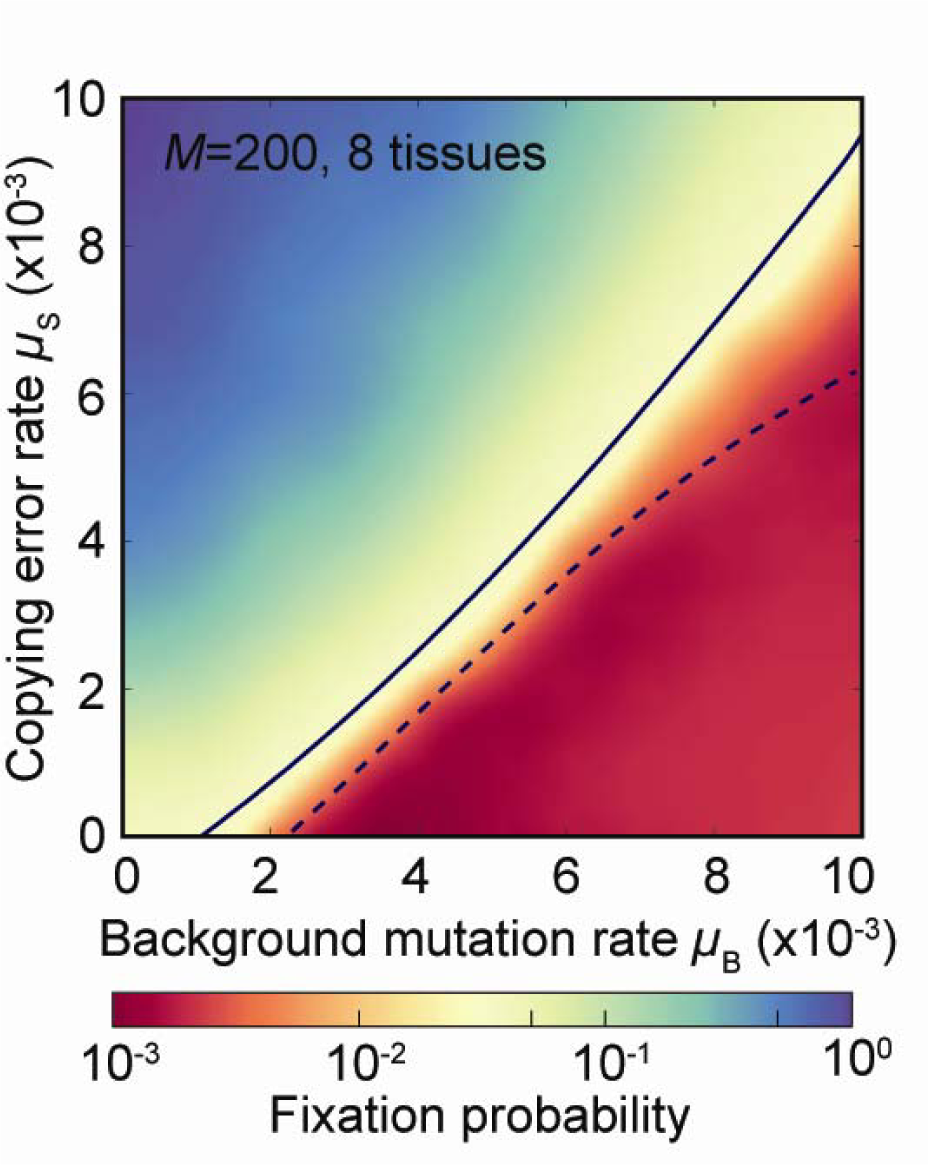
Greater complexity does not favour germline sequestration. Heat map showing the fixation probability of an allele encoding early germline sequestration (introduced at a frequency of 0.05, see Methods) in an organism with 8 tissues and 200 mitochondria. Early germline sequestration is favoured by higher *µ*_S_ and lower *µ*_B_ (blue, top left). The allele is selected against in organisms with low *µ*_S_ and high *µ*_B_ (red, bottom right), instead favouring somatic gametogenesis. The solid line represents neutrality, and the dotted line gives neutrality for the case shown in Fig. 3a, with no tissues and *M* = 50. Increasing the level of complexity (more tissues and mitochondria) therefore does not favour early germline sequestration.

**Extended Data Fig. 2.**
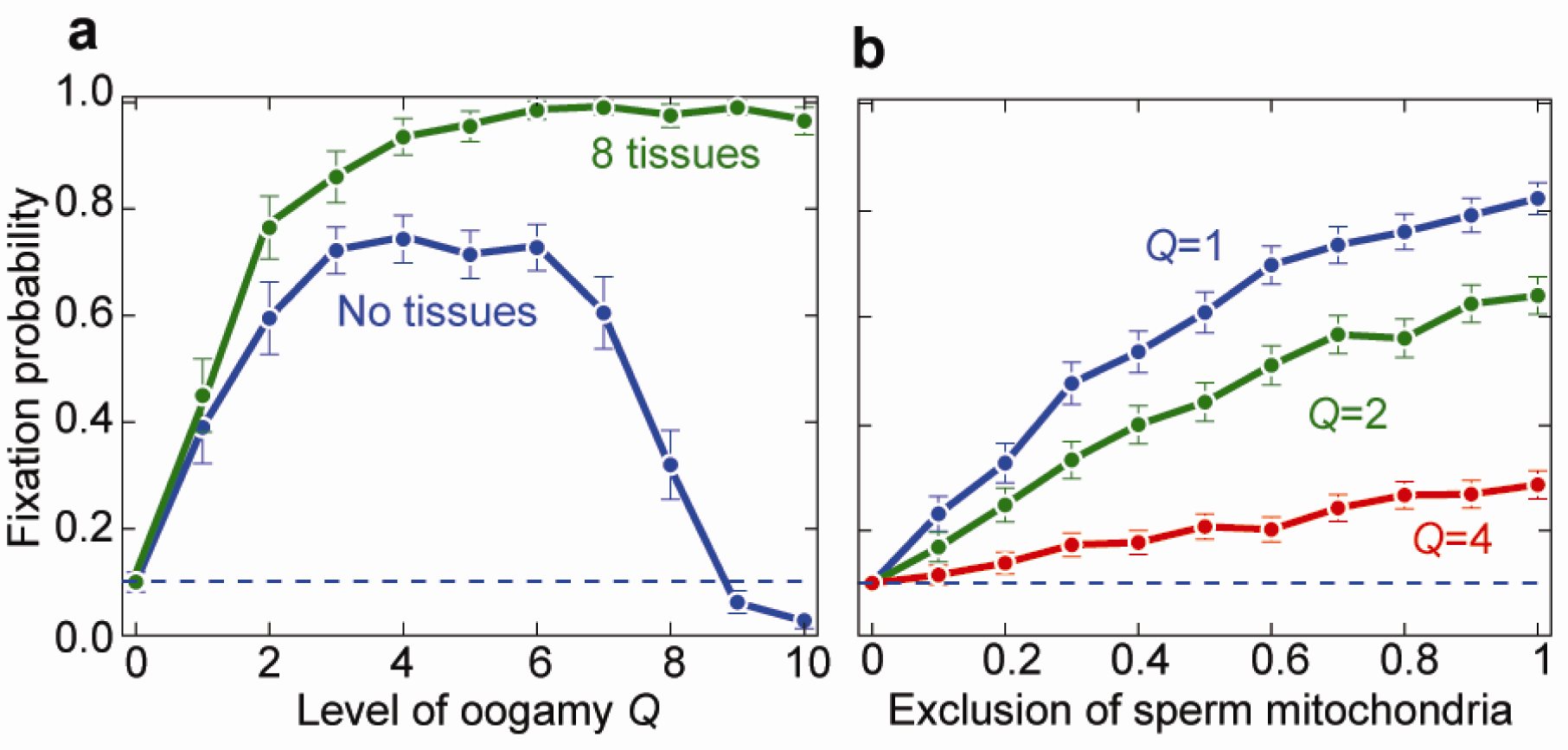
The evolution of anisogamy and strict uniparental inheritance is favoured in basal metazoans without tissues. **(a)** Oogamy (*Q*) is favoured even in organisms with no tissue differentiation if the starting point is biparental inheritance (BPI) of mitochondria. This is because oogamy produces a degree of partial uniparental inheritance (UPI) of mitochondria, which increases the variance between gametes and homogenizes the mitochondrial population within individual gametes. BPI is not an unreasonable starting point in basal metazoans as next generation sequencing shows that heteroplasmy and paternal leakage are not uncommon^50^; the disparate mechanisms that enforce UPI imply that strict UPI arose on multiple occasions and are subject to frequent turn-over^51^, hence some degree of BPI must be common over evolutionary time; and previous modelling work shows that UPI is less strictly enforced in organisms with low mitochondrial numbers and low mutation rates^52,53^, conditions that apply to basal metazoans such as sponges, which have very low, plant-like mitochondrial mutation rates^26-28,30^. Very high levels of oogamy (*Q*) are not favoured in the absence of tissues as this restricts segregational variance between gametes even with somatic gametogenesis, and there is no benefit to restricting early variance between tissues (which is why higher levels of *Q* are still favoured with multiple tissues). **(b)** The exclusion of male mitochondria from the zygote is favoured because that also increases the degree of UPI. The effect weakens with higher *Q* as the fractional contribution of sperm falls simply because there are fewer of them. The evolution of oogamy combined with exclusion of male mitochondria from the zygote is an alternative pathway to the evolution of strict UPI. We propose that this pathway could account for the evolution of anisogamy with stable UPI in basal metazoans. Parameter values *µ*_S_ = 0.001, *µ*_B_ = 0.005 and *M*= 50.

**Extended Data Fig. 3.**
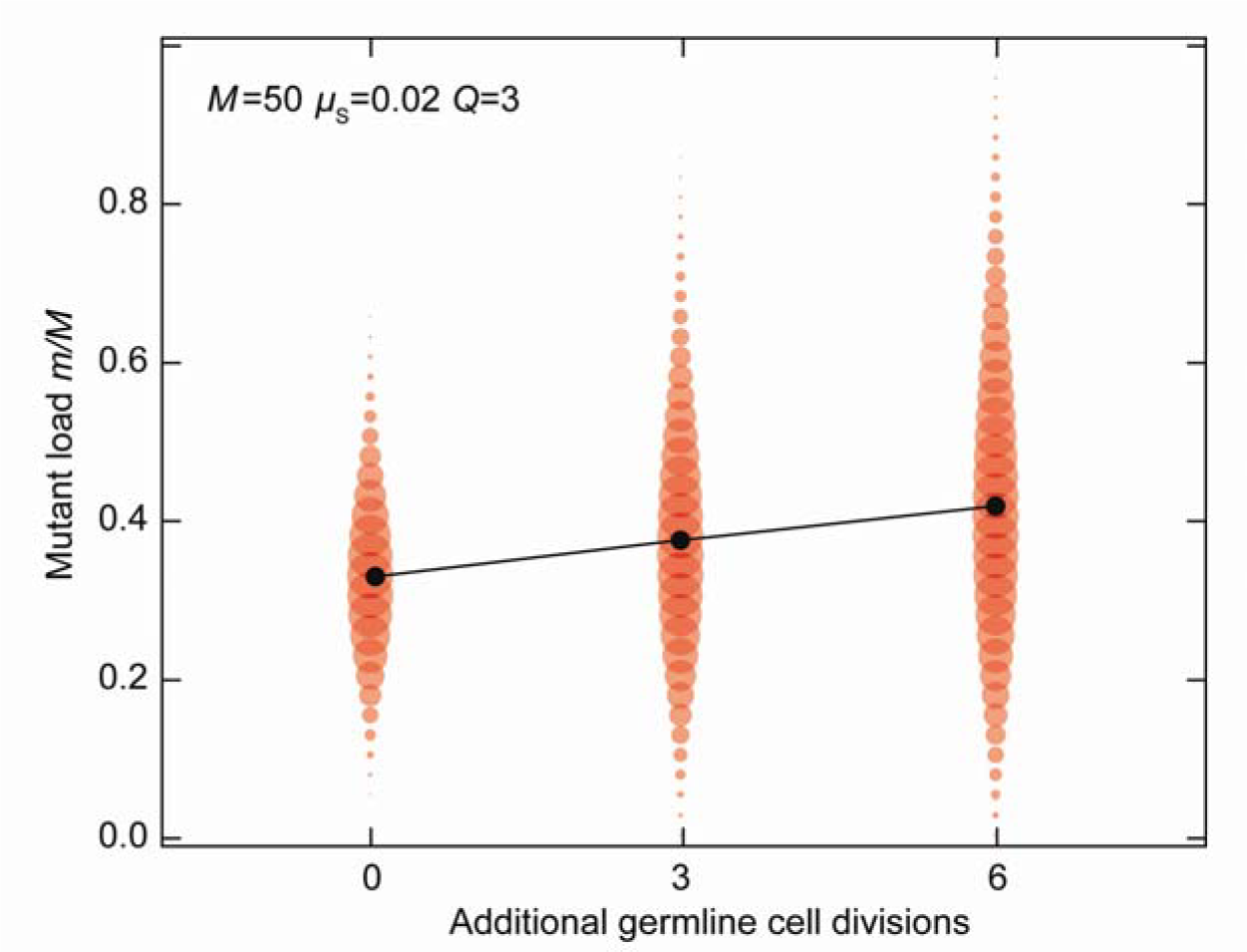
Restoration of germline segregational variance after oogamy. Additional cell divisions in the female germline restore the loss of segregational variance as a result of oogamy *Q*. In this example, *Q* = 3 and there is a high error rate (*µ*_S_ = 0.02). Additional cell divisions generate a greater mutational load, but also considerable segregational variance. Under these conditions, the balance between mutation accumulation and variance favours an intermediate germline (*N_G_* = 6, i.e. 3 additional germline cell divisions, once 3 rounds of early cell division returns cell content to *M* mitochondria) rather than an early germline (*N_G_* = 3, no additional germline cell divisions) or a late germline (*N_G_* = 9 or more). Random atresia at *N_G_* = 6 retains the same level of variance in the surviving cells between a smaller number of gametes. Other parameter values *µ*_B_ = 0.005, *M* = 50.

**Extended Data Fig. 4.**
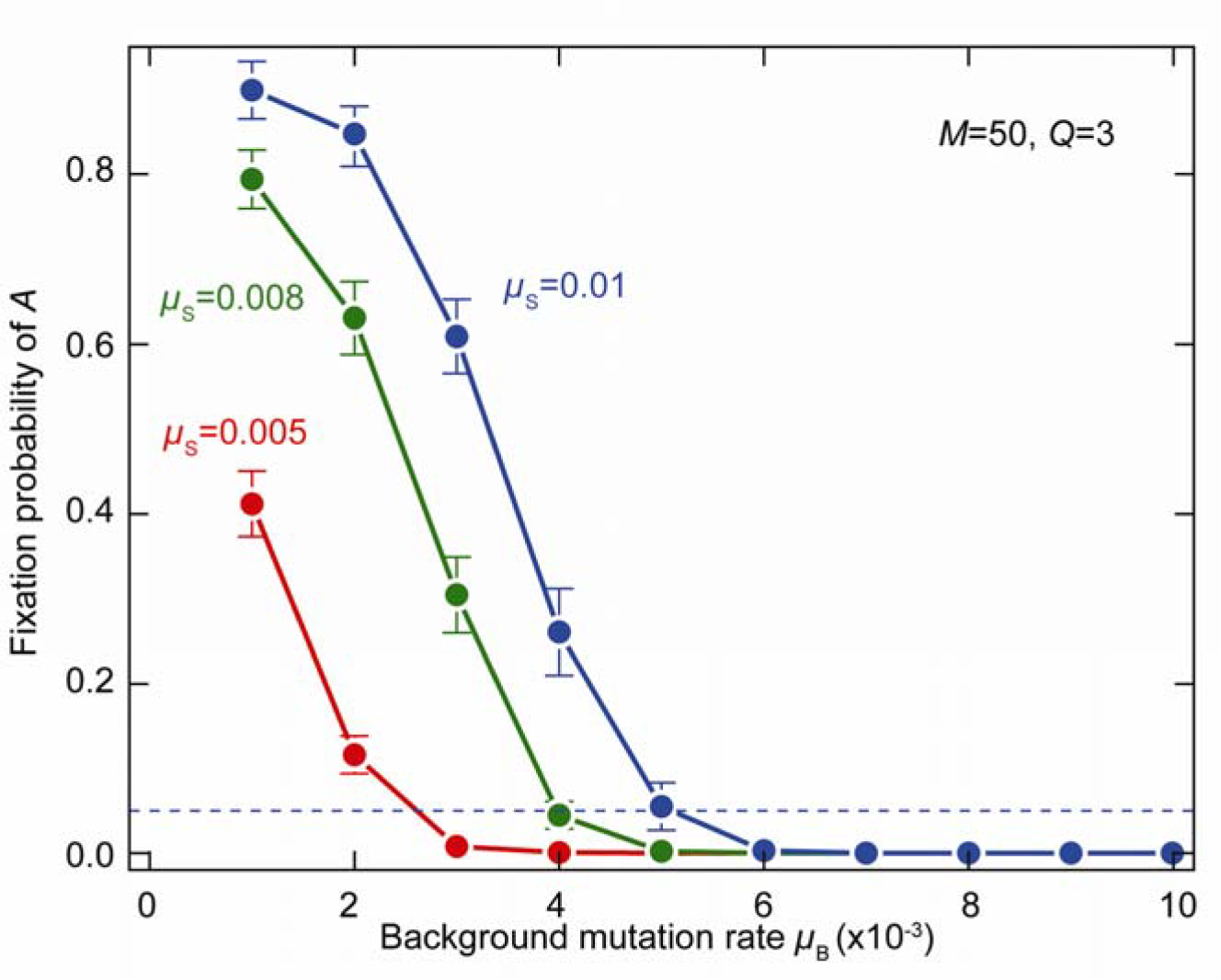
Restricting background mutations favours germline fixation. The evolution of early germline sequestration (*N_G_* = 3) and oogamy (*Q* = 3) is favoured by low background mutation rates (*µ*_B_). The allele *A* specifying early germline sequestration is more likely to be fixed at higher values of *µ*_S_, as high *µ*_S_ favours restricting the number of rounds of cell division, but at the cost of restricting variance. Germline fixation also depends on the background mutation rate (*µ*_B_). Germline sequestration does not fix if *µ*_B_ > 6 × 10^−3^, but fixes readily at lower *µ*_B_, even when *µ*_S_ is low. Adaptations that favour low *µ*_B_ are characteristic of bilaterians germlines, including sequestration of oocytes in an internal ovary, repression of transcription and translation, and suppression of mitochondrial respiration^14^.

## Methods

### Segregational drift generates variance in mitochondrial mutation load

Here we review the mitochondrial dynamics at the cell level, showing that random segregation at every cell division generates variance in the mutational load (Figures 2a-d and 4a-b). We start with a cell containing *M* mitochondria in total, out of which *m*_0_ are mutant. The initial state of an infinite population is then represented by a state vector **p**^(0)^, with all of its *M* + 1 entries set to zero, except for the *m*_0_-th, which is set to one, i.e. 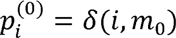. A single cell cycle consists of mitochondrial mutation, replication and cell division. Mitochondrial mutation is a Bernoulli trial with success probability *μ* while segregation is modelled as a simple random sampling without replacement. After cell divisions the population state vector becomes

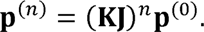

Here **J** is the (*M* + 1) × (*M* + 1) matrix with elements representing the transition probabilities due to mutation,

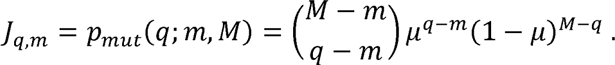

Similarly, **K** is the matrix of same dimensions representing the transformation due to random sampling,

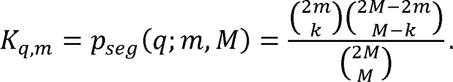

In Figure 2 we show distributions **p**^(*n*)^ for *n* = 0 … 10.

Variance in the mutant load after *n* cell divisions can be expressed analytically in the illustrative case of *μ* = 0, where only segregational drift is accounted for. Let *X_n_* be a random variable denoting the number of mutant mitochondria within a cell in the developing tissue or embryo after *n* cell divisions, and *x_n_* its actual realization. For sampling from the hypergeometric probability distribution, the population mean equals the initial number of mutants, E(*X_n_*) = *x*_0_, while the variance can be decomposed as

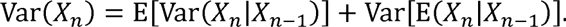

For sampling without replacement from the hypergeometric probability distribution, the conditional variance is

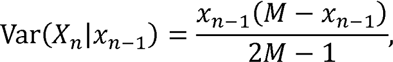

and so

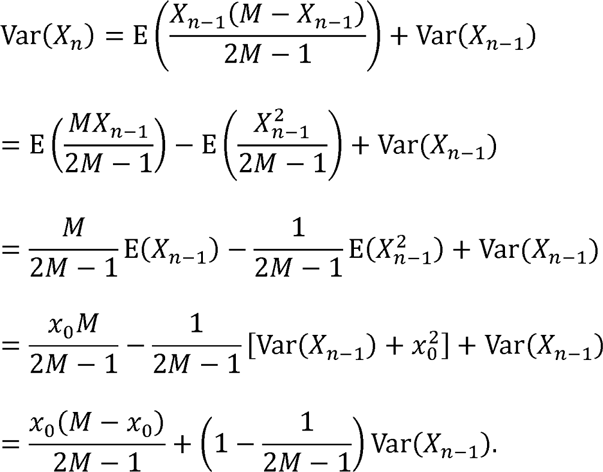

Here *x*_0_ is the initial number of mutants within a cell. This is a recurrence relation of the form *h_n_* = *Hh*_*n*−1_ with *h_n_* = *Var*(*X_n_*) − *x*_0_(*M* − *x*_0_) and 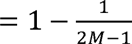. With the boundary condition Var(*X*_0_), *h*_0_ = −*x*_0_(*M* − *x*_0_), the solution is

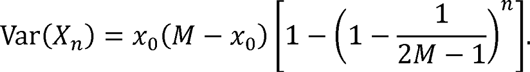

Variance in the mutant frequency 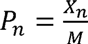 is then

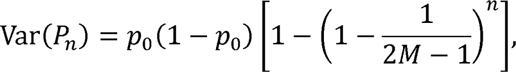

which corresponds to Figure 4b.

In oogamous matings, the zygote contains 2^*Q*^*M* mitochondria and undergoes 3 divisions without mitochondrial replication. Changes in variance follow a similar trend, but this time the recurrence relation is

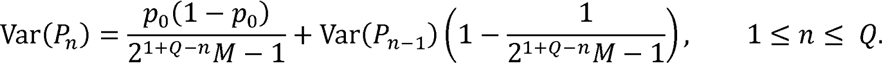

The recurrence relation can be used directly to produce Figure 5b For a large initial number of mitochondria 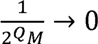 and the approximate solution is

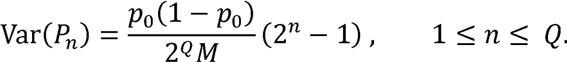

### Finite-population simulation model for the evolution of germline

To study the evolutionary dynamics of alleles responsible for germline sequestration we developed a multi-level agent-based model, implemented as a set of simulation routines in ANSI/ISO C++ (https://github.com/ArunasRadzvilavicius/GermlineEvolution). The model population consists of *N* = 500 multicellular individuals, with equal numbers of males and females. A single individual is represented as an object containing a variable number of cells, either undifferentiated or grouped into tissues while each cell is an object containing the number of mutant and wild type mitochondria. Generations are discrete and non-overlapping. Organism development proceeds by iterating through the population and modifying these objects according to a set of predefined rules. The population is initialized in a random state, and the simulation proceeds as follows:

1. Start the generation with a population of *N* zygotes;
2. For each organism in the population, repeat until development is complete:
  a. For every somatic cell within an organism:
    i. Apply background mitochondrial mutation as a Bernoulli event with the success probability *μ_B_*;
    ii. If the number of mitochondria is *M*, apply mitochondrial mutation as a singular Bernoulli trial with success probability *μ_s_*, then duplicate the mitochondrial population;
    iii. Partition mitochondria into two daughter cells by sampling without replacement, which in practice is implement by drawing a random number form the hypergeometric probability distribution;
  b. If the number of cell divisions equals *G*, set aside a single primordial germ cell by copying a randomly chosen stem cell, which will not undergo further mitotic divisions;
  c. If the number of cell divisions equals *T*, assign each of 2^*T*^ stem cells to its own somatic tissue;
  d. If the number of cell divisions equals *L*, stop the development. For each cell apply the background mitochondrial mutation *S* = 40 times independently, each time with probability *μ_B_*;
3. Apply selection by sampling *N*/2 males and *N*/2 females from the population with replacement, linearly weighted according to their mitochondrial fitness;
4. In female germ cells, repeat *Q* times:
  a. Apply mitochondrial mutation as a binomial event with probability *μ_s_*, duplicate the mitochondrial population
5. Complete gametogenesis by executing two meiotic cell divisions, with mitochondrial sampling without replacement; male gametes contain *M*/2 and eggs contain 2^*Q*^*M*/2 mitochondria;
6. Fuse random gamete pairs of opposite sexes, making sure that the zygote contains 2^*Q*^*M* mitochondria.

Gametes of opposite sexes meet at random and fuse. Parameter *v* controls the amount of mitochondria inherited from the father (paternal leakage). At fertilization (step 6), the zygote’s mitochondrial population of 2^*Q*^*M* is formed by sampling (1 − *v*)*M*/2 mitochondria from a sperm cell, and the rest from the oocyte.

### Evolution of the nuclear alleles and fixation statistics

Mating type, the number of germline cell divisions *G*, the log-size of the zygote *Q* and the degree of uniparental inheritance *v* are all traits of an organism controlled by a set of loci in the nuclear genome, which is assumed to be diploid. The mating type locus is heterogametic *ZW* in females, while the other mating types is homogametic *ZZ*.

We determine the selective advantage of an invading allele in a Monte Carlo simulation, by numerically calculating its fixation probability within a finite population. With the population at equilibrium, the invading allele is introduced at a low frequency *f*_0_ = 0.05 (or 0.1), and its fate is tracked until either fixation or extinction. This requires 10^3^ − 10^4^ repetitions of the calculation depending on the mutation rates and the associated levels of noise. This pattern is assessed relative to the fixation probability of a neutral allele, which simply equals its initial frequency *f*_0_. The error bars in figures represent the 95% confidence interval of the binomial proportion via the Gaussian approximation, that is 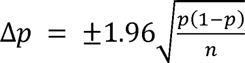, where *n* is the number of trials and *p* is the measured rate of fixation. An allele is deemed to be evolutionarily advantageous if its fixation probability exceeds the chance of fixation of the neutral allele.

The same general procedure is applied to determine the fate of modifiers coding for the number of germline cell divisions, the level of mitochondrial oogamy or mitochondrial exclusion. The germ cell differentiation locus is expressed in both sexes, and is assumed to be autosomal, with the invading allele assumed to be dominant. We also examined other dominance states, but the main conclusions of this work remained unaffected. The other two loci are *W* linked and unaffected by dominance considerations.

